# Uncertainty in population estimates: a meta-analysis for petrels

**DOI:** 10.1101/2021.04.07.438872

**Authors:** Jeremy P. Bird, Bradley K. Woodworth, Richard A. Fuller, Justine D. S. Shaw

## Abstract

1. Population estimates are commonly generated and used in conservation science. All estimates carry inherent uncertainty, but little attention has been given to when and how this uncertainty limits their use. This requires an understanding of the specific purposes for which population estimates are intended, an assessment of the level of uncertainty each purpose can tolerate, and information on current uncertainty.
2. We conducted a review and meta-analysis for a widespread group of seabirds, the petrels, to better understand how and why population estimates are being used. Globally petrels are highly threatened, and aspects of their ecology make them difficult to survey, introducing high levels of uncertainty into population estimates.
3. We found that by far the most common intended use of population estimates was to inform status and trend assessments, while less common uses were trialling methods to improve estimates, and assessing threat impacts and conservation outcomes.
4. The mean coefficient of variation for published estimates was 0.17 (SD = 0.14), with no evidence that uncertainty has been reduced through time. As a consequence of this high uncertainty, when we simulated declines equivalent to thresholds commonly used to trigger management, only 5% of studies could detect significant differences between population estimates collected 10 years apart for populations declining at a rate of 30% over three generations.
5. Reporting of uncertainty was variable with no dispersion statistics reported with 38% of population estimates and most not reporting key underlying parameters: nest numbers/density and nest occupancy. We also found no correlation between population estimates and either island size, body size or species threat status—potential predictors of uncertainty.
6. *Synthesis and applications*—Key recommendations for managers are to be mindful of uncertainty in past population estimates if aiming to collect contemporary estimates for comparison, to report uncertainty clearly for new estimates, and to give careful consideration to whether a proposed estimate is likely to achieve the requisite level of certainty for the investment in its generation to be warranted. We recommend a practitioner-based Value of Information assessment to confirm where there is value in reducing uncertainty.

## INTRODUCTION

Population estimates are a foundation of conservation science, variously used to characterise the changing status of populations, manage harvests, assess threats to biodiversity, choose between and evaluate outcomes of conservation actions, and to communicate conservation messages (Jones et al., 2016; Nichols et al., 2007; Pressey et al., 2007; Rodrigues et al., 2006). Inherent in all population estimates is a level of uncertainty. Logically, the lower the uncertainty, the easier it is for managers to use an estimate to make an informed decision, but not all uncertainties are important to resolve (Runge et al., 2011). Uncertainty in a population estimate matters only where a course of action might change if that uncertainty is reduced. Given the trade-off between spending money on surveying biodiversity and spending it on management there is value in understanding the return on investment from estimating population sizes, and the benefit of reducing uncertainty of those estimates (Possingham et al., 2012; Runge et al., 2011).

Many biodiversity censusing and monitoring studies do not clearly state the purpose for their monitoring (Possingham et al., 2012). While conservation interventions may report their proximal goals, many fail to outline their ultimate objectives (Bird et al., 2019; Mackenzie & Royle, 2005). For example, the proximal goal of poison-baiting an island may be to eradicate an invasive predator, but its ultimate goal of precipitating an increase in the population of a threatened species is not always stated or given a quantitative target. In such cases it is not possible to determine whether reducing uncertainty around a population estimate is worthwhile. If there is no direct mention of an outcome that will be informed by a population estimate, it begs the question is there any need to make that estimate? In cases where the purpose for generating a population estimate is clearly articulated, the cost of the estimate can be weighed against the benefit of reducing uncertainty around it (Canessa et al., 2015). While population estimates are widely reported and advocated, and are presumably attracting significant conservation resources, there has been little analysis of why population estimates are being generated, how much uncertainty they carry, and whether uncertainty in those estimates might hinder use for their stated management purpose.

Here we investigate uncertainty in population estimates for an entire taxonomic group of seabirds, the petrels (families *Procellariidae, Hydrobatidae* and *Oceanitidae*). Many studies have flagged uncertainty in population estimates as an issue, highlighting potential weaknesses of individual survey methods, and challenges in particular study systems (e.g. Arneill et al., 2019; Hatch, 2003; Sutherland and Dann, 2012). While these case studies hint at a potentially widespread issue, by reviewing the entire group we can assess trends in measurement and reporting of uncertainty and provide guidance on how it might best be tackled.

Understanding the size and trends of petrel populations is important ecologically and commercially because of the ecological function petrels play interacting with island-ocean food webs, fisheries and as a harvestable resource (Danckwerts et al., 2014; Graham et al., 2018; Newman et al., 2009; Otero et al., 2018). It is also important for conservation as 52 of 124 species (42%) are threatened with extinction (Rodríguez et al., 2019). Petrels nest almost exclusively on remote islands where monitoring is rare (Warham, 1996). Transport is a major component of costs, so infrequent but large-scale surveys tend to be more prevalent than frequent small-scale monitoring visits (Buxton et al., 2016; Rodríguez et al., 2019). As a result, changes are often inferred from a small number of measures, such as species population estimates, obtained many years apart (e.g. see Brooke, Bonnaud, Dilley, Flint, Holmes, Jones, Provost, Rocamora, Ryan, Surman, et al., 2018). Apparent changes detected from repeated population estimates have catalysed invasive species eradication programmes (Brooke, Bonnaud, Dilley, Flint, Holmes, Jones, Provost, Rocamora, Ryan, & Surman, 2018), informed fisheries and harvest management including influencing policy and international agreements (Agreement on the Conservation of Albatrosses and Petrels, 2001; Newman et al., 2009; Richard & Abraham, 2013), been used to assess the outcomes of conservation interventions and enhance future ones (Bourgeois et al., 2013; Parker et al., 2015), and to inform Red List status assessments (Rodríguez et al., 2019). However, high uncertainty in petrel population estimates can hamper their ability to achieve and unambiguously inform these objectives. For example, Cory’s Shearwaters *Calonectris borealis* were surveyed on the island of Corvo in the Azores to assess population trajectories in response to known threats, but the study concluded that uncertainty in the final estimate was too great to detect moderate changes in population size (Oppel et al., 2014). Flesh-footed Shearwaters *Ardenna carneipes* on Lord Howe Island are prone to plastic ingestion but conclusions about the population-level impact of this potential threat are clouded by changes in equipment and survey design over successive attempts to estimate the population size (see Carlile et al., 2019; Lavers et al., 2019).

For a considerable majority of petrels obtaining accurate estimates is particularly difficult because nests cannot be counted directly. Instead they are hidden in crevices and underground burrows distributed heterogeneously across challenging terrain, and birds only return to or leave from their burrows at night (Rayner et al., 2007; Schumann et al., 2013; Warham, 1996). Uncertainty around petrel population estimates is inflated because variance around both the estimated number of burrows and the proportion of burrows occupied by breeding pairs is propagated when the two metrics are combined (Whitehead et al., 2014). Given the inherent uncertainties in petrel population estimates, and the debate around spending limited funds on undertaking conservation actions or gathering evidence to inform future actions (Brooke, Bonnaud, Dilley, Flint, Holmes, Jones, Provost, Rocamora, Ryan, & Surman, 2018), it is important to know whether, and how much, the level of uncertainty in estimates matters.

We undertook a literature review and meta-analysis to: (i) understand the motivations for estimating population sizes for petrels, (ii) assess uncertainty in population estimates and correlates thereof; and (iii) evaluate the implications of uncertainty given the prevalent but potentially flawed use of population estimates for detecting population change. We examine whether uncertainty in population estimates has improved over time and assess whether there are clear steps that can be taken to minimise uncertainty in future estimates to improve management.

## METHODS

### Approach

Our study had two parts: (a) a review of published literature reporting estimates of petrel population sizes on islands to explore motivations for generating estimates, and (b) a meta-analysis of uncertainty in published population estimates. Throughout we restrict our review to petrels that nest underground. In our meta-analysis we conducted a simulation to understand the power for published estimates to detect population change given their reported uncertainty. To examine drivers of uncertainty, we investigated first whether the number of estimates being published has changed over time and if uncertainty is related to publishing trends, and then whether uncertainty can be explained by variables between estimates.

### Literature review

We searched the Web of Science bibliographic index on 20 January 2020 (see Appendix S1 in the Supporting Information). Our structured search returned 900 studies. We screened all titles and abstracts for papers potentially reporting population or colony estimates for any of our target species (see Table S1 in the Supporting Information) from one or more islands, and then assessed full text articles, retaining all papers that reported quantitative estimates (Figure 1). After full screening, 68 studies were selected for review. All studies in the literature review and meta-analysis are available from the Data Availability Statement.

**Figure 1:**
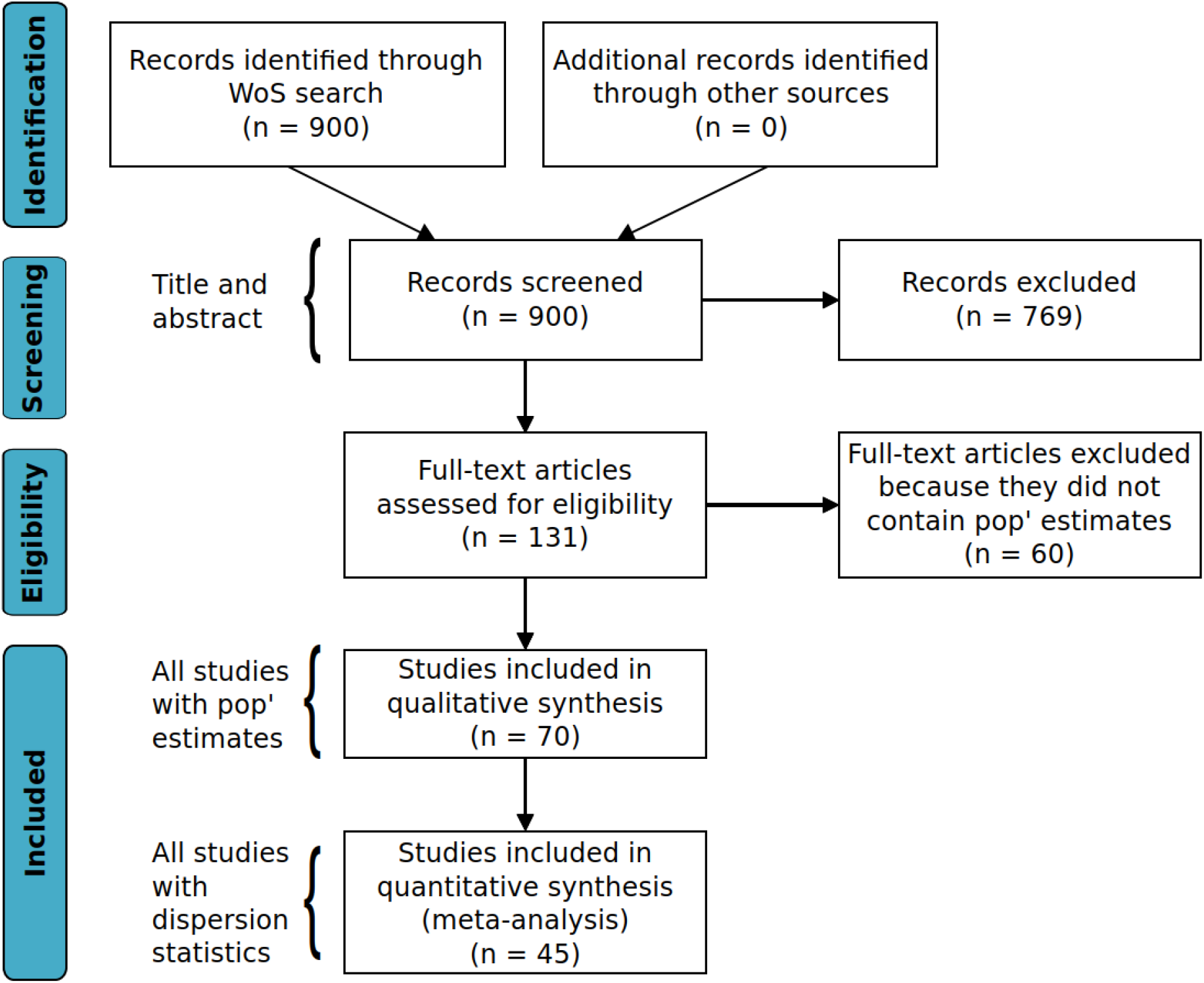
PRISMA flowchart adapted from Moher et al. (2009) documenting the search and screening steps in our review.

Before conducting our review, we gave prior consideration to the main reasons people might estimate petrel population sizes. For each reason we proposed a level of requisite certainty that estimates must attain to be fit-for-purpose (Table 1). During the review, for each paper containing a population estimate we recorded any mention of specific management actions the paper intended to inform and recorded categorically all reasons explicitly mentioned as motivations for the survey/population estimate.

**Table 1:**
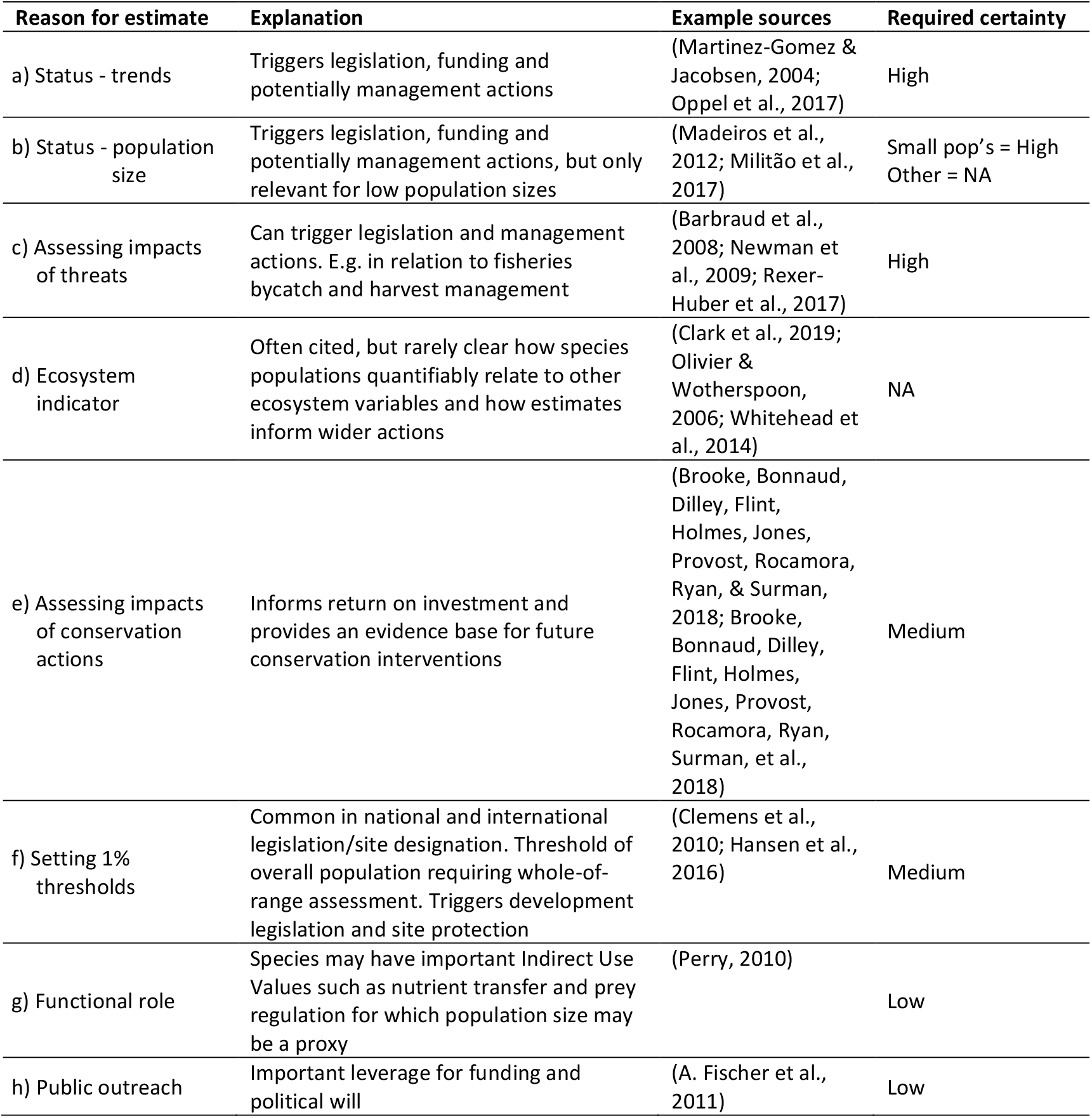
Reasons to estimate petrel populations and required certainty identified *a priori*. We ranked required certainty based on our interpretation of the financial, societal and conservation implications of incorrect data.

### Meta-analysis

We extracted published population estimates reported in each paper. Most reported a mean (*μ*), but where only minima or maxima were reported we used this as the estimate, and where only minima and maxima were reported we used their average as the estimate. If available, we extracted reported 95% confidence intervals (CI). If CI was not reported, but standard error 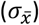 was we estimated: 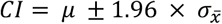. To compare reported uncertainty between studies we calculated the coefficient of variation (CV) as 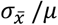.

#### Simulated power analysis

We used simple simulations to assess the power to detect population declines from published population estimates and their uncertainty levels. Using the mean population estimate as the initial population size (*N*_*1*_) for each case study, we first generated a sequence of annual true population sizes (*N*_*t*_) for declines of 30%, 50%, and 80% over 3 generations (*t*_*max*_)(following Bird et al., 2020; IUCN, 2012). These rates of decline, measured entirely in the past, in the past and future, or entirely in the future are thresholds for listing species as Vulnerable, Endangered and Critically Endangered on the IUCN Red List and correspond to rapid, very rapid and extremely rapid population declines (BirdLife International, 2020). Population declines were generated based upon an exponential rate of population change (*N*_*t*_ = *N*_*1*_ * *e*^*rt*^, where *r* is the exponential rate of change) with no stochasticity. For each time-series, we then simulated 20,000 Monte Carlo (MC) samples at each time step from a normal distribution with mean *N*_*t*_ and variance approximated from the published confidence intervals. For these simulations, we assumed a fixed coefficient of variation, such that variance scaled with the mean. For each MC sample, we then calculated the proportional population change between two samples (*N*_*1*_ and *N*_*t*_) as *N*_*t*_*/N*_*1*_ for *t* in *1:t*_*max*_, where *N*_*t*_*/N*_*1*_ = 1 indicated no change and *N*_*t*_*/N*_*1*_ < 1 signalled a decline. Ratios were summarized across MC samples by the mean and 95% interval (2.5%, 97.5% quantiles). We considered ratios with upper 95% intervals < 1 indicative of significant population decline (see the Data Availability Statement for a link to all data and code).

#### Publishing trends and drivers of uncertainty

A number of sources contribute to uncertainty in petrel population estimates (Table S1, Supporting Information). A common challenge is that estimates are often derived from extrapolated burrow counts and sampled burrow occupancy, thus propagating uncertainty from multiple sources. To unpick their different contributions, we extracted available information on dispersion in reported burrow counts/density estimates and occupancy statistics. Population estimates were then grouped according to those that reported burrow and occupancy statistics and those that did not. Having derived the CV for each estimate (see above) we compared the means of these intervals between groups using t-tests, group trends in the size of normalised confidence intervals through time were assessed with linear regression, and trends in the proportion of studies reporting different variances were tested with Chi-squared tests of proportions. To allow analysis and identification of potential correlates of uncertainty we extracted additional variables reported with the published population estimates: the year of study/survey, body size and Red List status of the surveyed species, size of the island surveyed, and the survey methods used (Table 2).

**Table 2:**
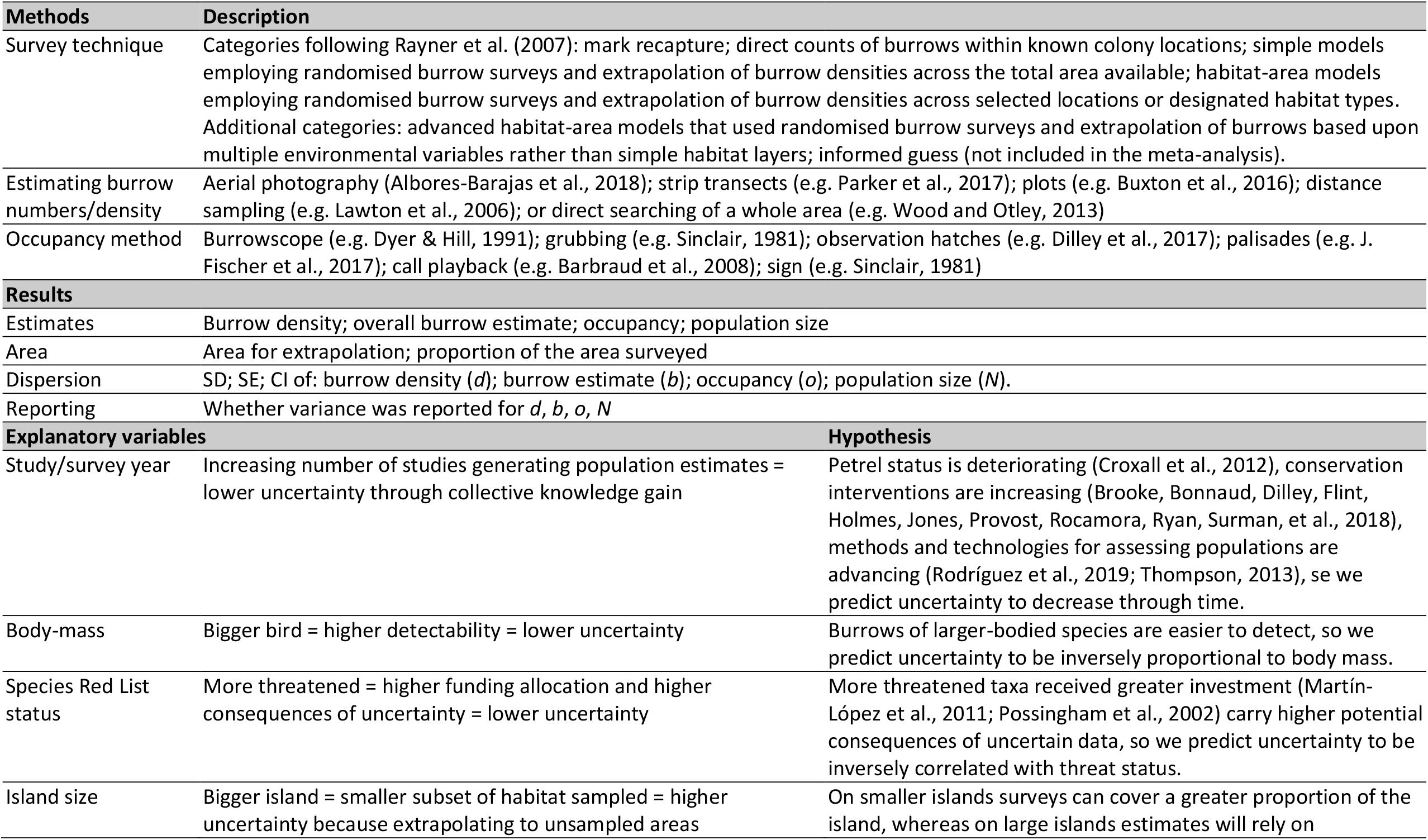
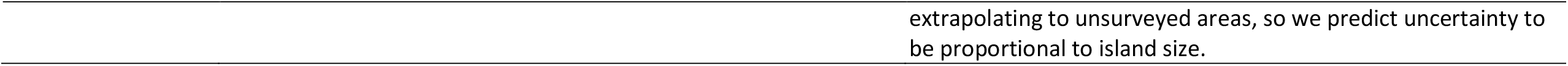
Variables recorded during literature review

We tested for correlations between explanatory variables and uncertainty (Table 2). Although these variables are beyond the control of managers it may be possible to use them to predict whether future surveys are more or less likely to achieve a reasonable level of uncertainty, or to identify where further research might be focussed to reduce uncertainty (e.g. improving survey methods for large islands or small-bodied species).

We fitted a global linear model including all explanatory variables and the survey technique (Table 2) as fixed effects, study as a random effect, and normalized confidence intervals as the response to identify predictors of uncertainty. We then fitted simpler models consisting of all combinations of subsets of the explanatory variables and compared models using Akaike’s Information Criterion (AIC). Candidate models were selected following Richards (2015). All models within 6 AIC of the preferred model (lowest AIC) were considered but a model was only included with the preferred model if there were no more parsimonious models with lower AIC. We also plotted bivariate relationships between normalised confidence limits and each explanatory variable and assessed their explanatory power using adjusted *R*^*2*^ (continuous variables) or ANOVA (categorical variables). All analyses and visualisation was performed in ‘R’ version 4.0.1 using base functions, the ‘tidyverse’ and lme4 packages (Bates et al., 2015; R Core Team, 2020; Wickham et al., 2019).

## RESULTS

### Literature review

We identified 68 studies that reported 170 population estimates in total. Some studies reported more than one motivation for generating a population estimate. The motivations reported had all been identified *a priori* as reasons for generating estimates (Table 1), except for studies focussed on methods to improve estimates (*n* = 14). Assessing species status and trends by providing a baseline or comparison figure for trend estimates was the reported motivation for 51 (75%) of 68 studies (Figure 2). For 87 (67%) of the 129 estimates published in these 51 studies, a historic estimate for the same species and site was referenced, but in relation to only 47 estimates (under half) were these historic data used to assess population change. In the remaining 48 cases the paper reported that methodological differences precluded comparison between historic and contemporary estimates. Of the 42 cases where a comparison was made, on only one occasion was uncertainty around the reported change quantified.

**Figure 2:**
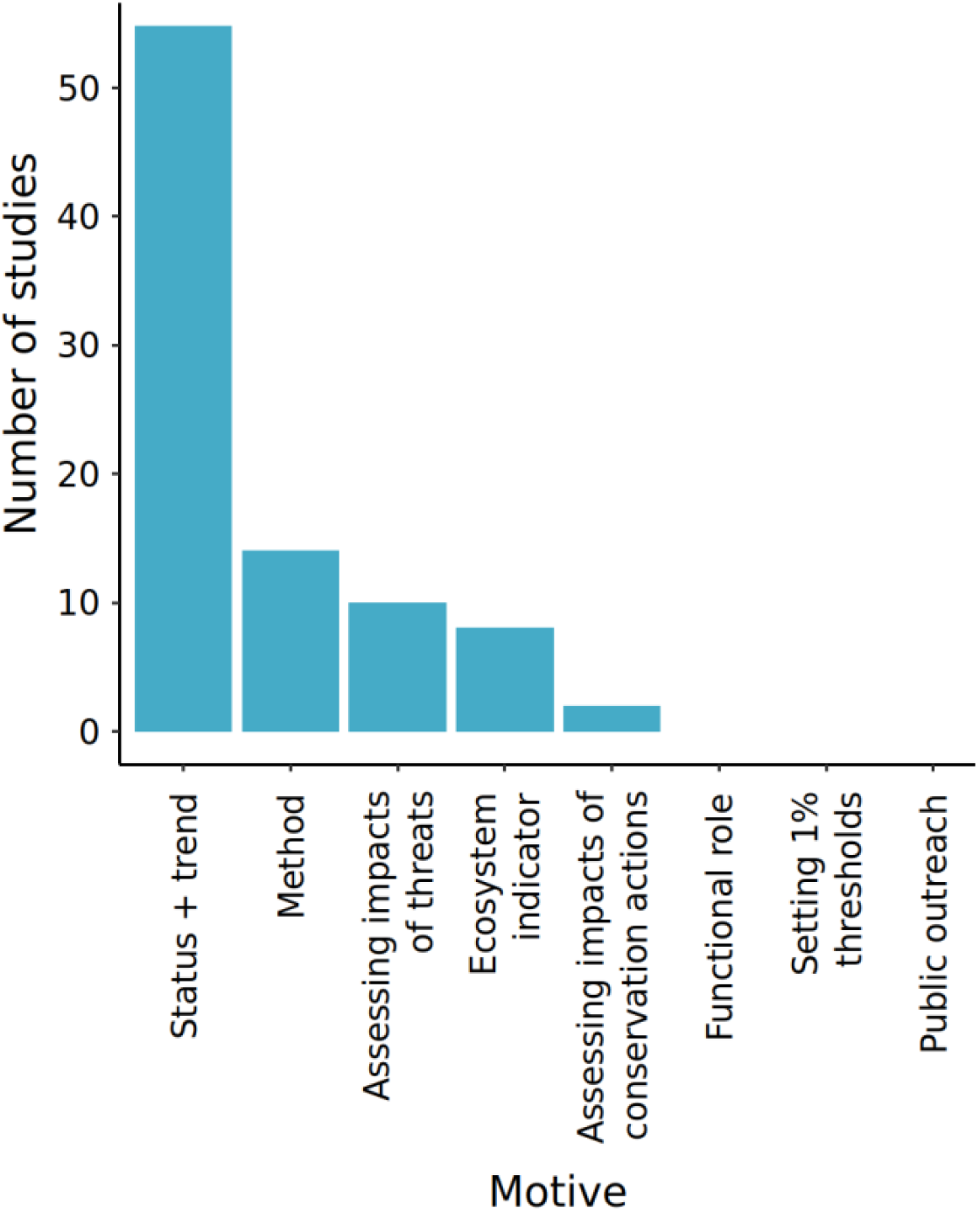
Motivations stated for estimating petrel population sizes from 68 studies. For detailed description of the motivations see Table 1. ‘Method’ was not predicted in Table 1 and reflects those studies for which the main motivation was to trial a method and test its utility for population estimation.

Four studies containing 12 estimates did not mention any reason for generating the estimates, and no studies reported collecting estimates to inform 1% population thresholds, to assess whether functional roles are being filled, or specifically for public outreach.

### Meta-analysis

Quantitative meta-analyses were restricted to 106 estimates from 44 studies that either had a confidence interval or reported dispersion statistics which we used to approximate a confidence interval. The mean CV of these estimates was 0.17 (SD = 0.14).

#### Simulated power analysis

For each of 106 population estimates with reported or approximated confidence intervals, we determined the minimum time between samples required for a statistically significant difference in population size to be detected for simulated declines of 30%, 50%, and 80% over 3 generations (Figure S1, Supporting Information). This is equivalent to measuring the minimum decline between samples that can be detected. We found that if populations were re-surveyed after 10 years, only 52% of studies could detect a change in a population declining at an annual rate consistent with 80% decline over three generations, 20% could detect changes in a population declining at an annual rate consistent with 50% decline over three generations, and just 5% of studies could detect changes in a population declining at an annual rate consistent with 30% decline over three generations (Figure 3).

**Figure 3:**
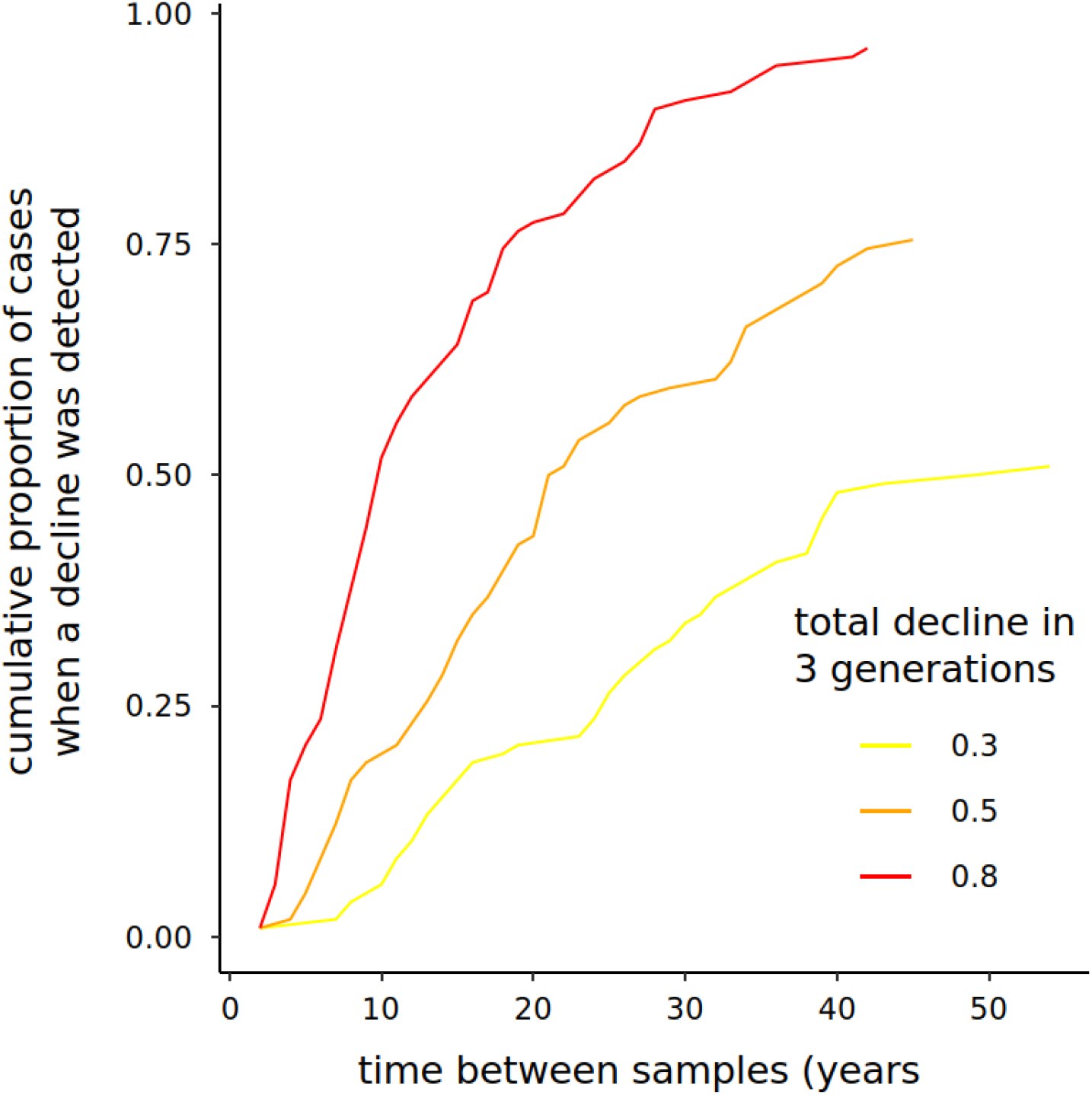
Based upon reported uncertainty, the proportion of studies able to detect significant population change after different lengths of time between repeat surveys given simulated rates of population declines equivalent to 30%, 50% and 80% over three generations (sensu IUCN 2020).

#### Publishing trends and correlates of uncertainty

The number of studies publishing population estimates (Figure S2, Supporting Information) and the proportion of them that also report uncertainty have both increased over time (Figure 4). Surprisingly, the magnitude of uncertainty in overall population estimates has also increased over time. We found no evidence that the proportion of studies that report key underlying parameters of most estimates, burrow numbers and burrow occupancy, has increased (Figure 4 and Figure S3 in Supporting Information). Reporting of both parameters is low. Of those studies that do report burrow numbers and burrow occupancy 76% and 33% respectively report uncertainty around these parameters and there is no evidence that these proportions have increased. Overall, uncertainty was significantly higher when estimating burrow numbers than burrow occupancy (Figure 4). We found some evidence that inspection hatches result in lower uncertainty than playback when estimating burrow occupancy, and that transects resulted in lower uncertainty than plots when estimating burrow numbers, but there were no significant differences between other methods reported (Figures S4 and S5, and Tables S2 and S3 in Supporting Information). There do not appear to be clear differences between species and genera among the taxa covered by our review (Figure S6 in Supporting Information).

**Figure 4:**
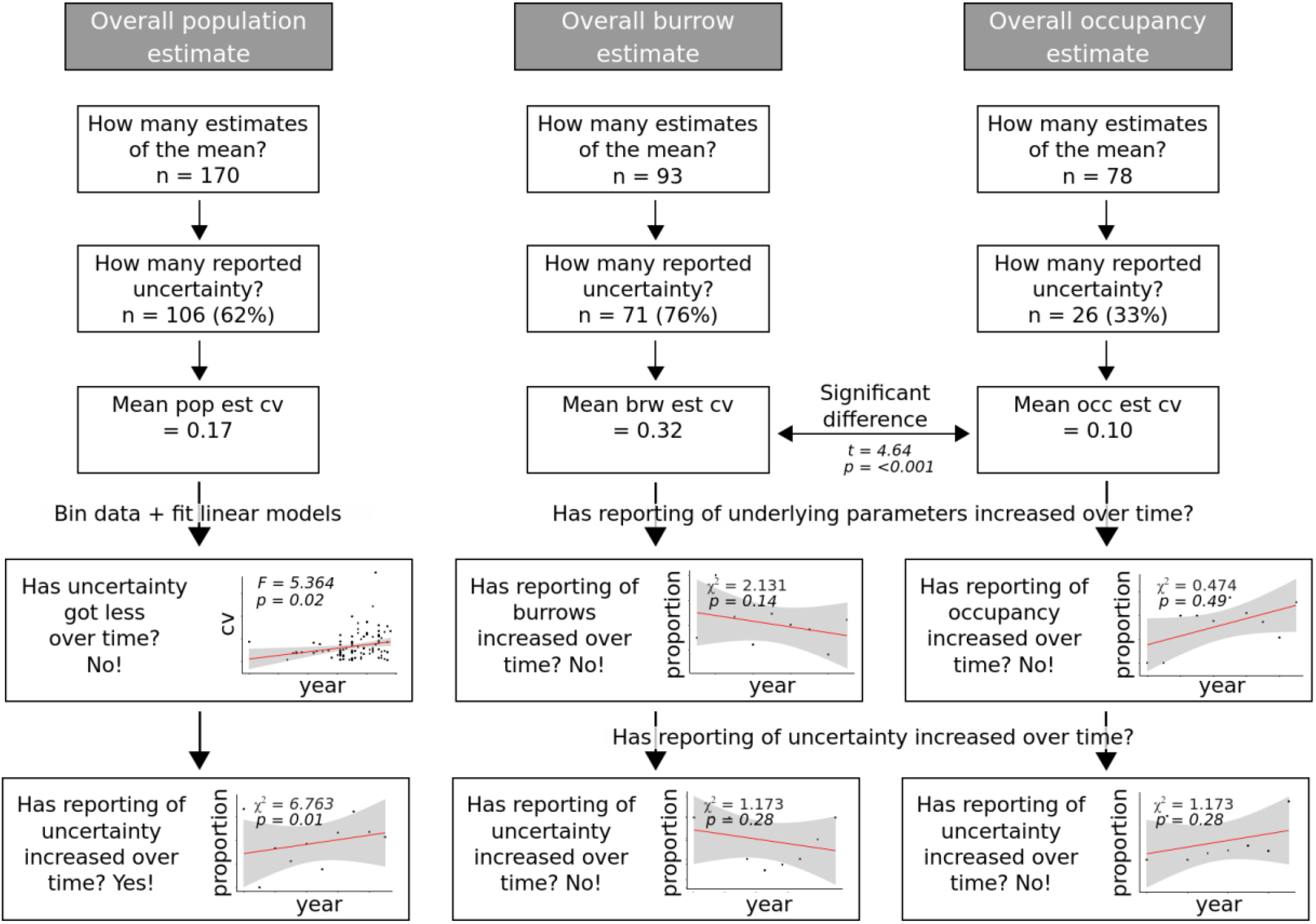
Reporting of petrel population estimates and the two key parameters that inform them, burrow numbers and burrow occupancy. Trends in reporting and the magnitude of uncertainty for all three parameters are included.

Of all candidate models from our multi-variate analysis an intercept only model, i.e. the mean CV with no explanatory variables, provided the best fit to the data (Table S4 in Supporting Information). Plotting the bivariate relationships supports this result with the only significant correlates of CV being population size and year of estimate, although the relationships were both weak (population size: *R*^*2*^ = 0.065; year of estimate: *R*^*2*^ = 0.04). We found no influence of species body mass, island size, Red List status or the survey method used (Figure 5 and Table S5 in Supporting Information). Thus, our analysis could not demonstrate that certainty in estimates is improving through time, or that estimates were more certain for larger, more easily detected species, or more threatened species (Figure 6).

**Figure 5:**
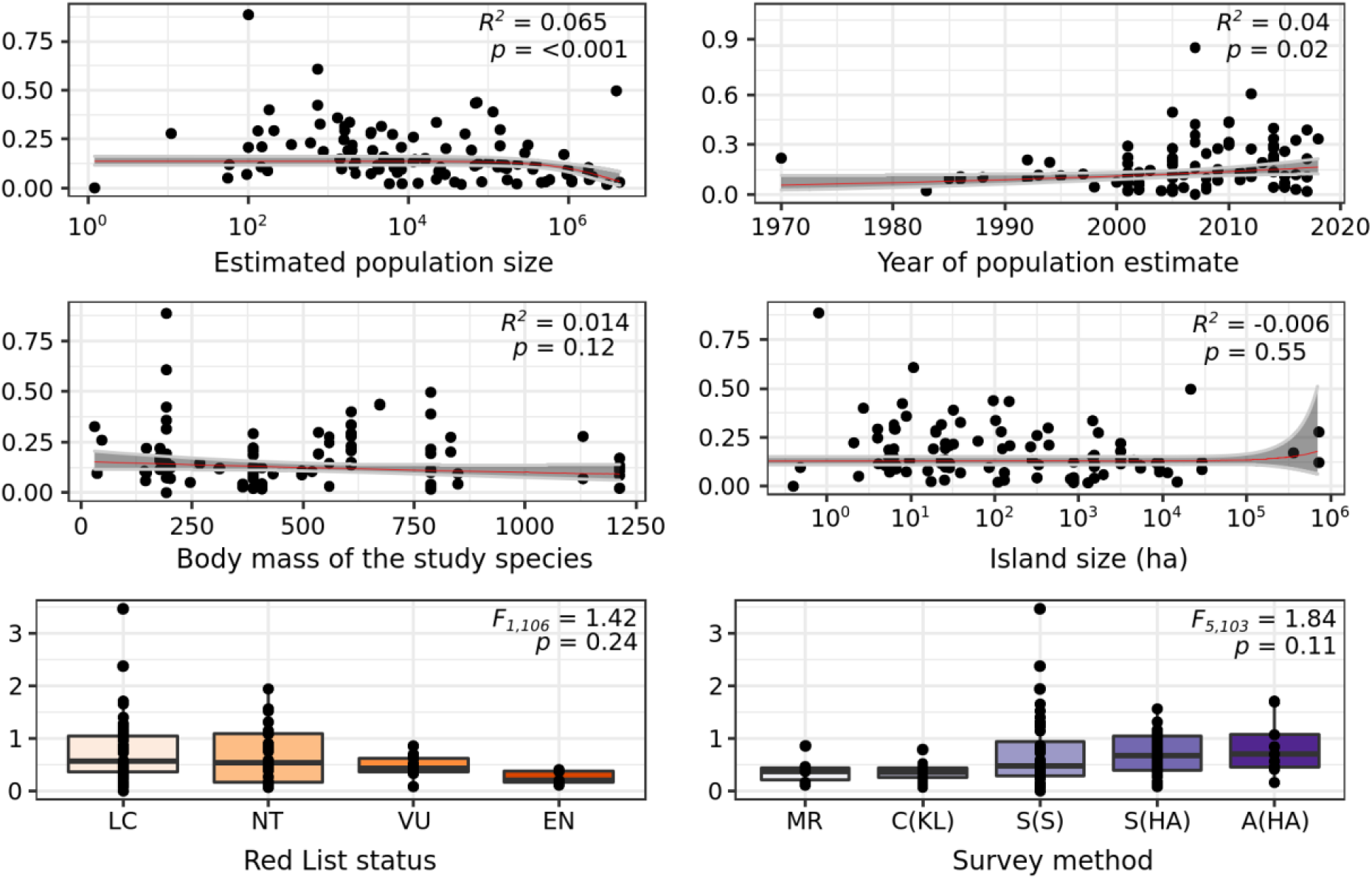
Uncertainty in petrel population estimates in relation to explanatory variables. Red List categories are: Least Concern (LC), Near Threatened (NT), Vulnerable (VU) and Endangered (EN). Survey methods are derived from Rayner et al. (2007): mark recapture (MR), counts of known colonies (C[KL]), simple extrapolation (S[S]), simple habitat area models (S[HA]), advanced habitat area models (A[HA]), informed estimates (IE).

## DISCUSSION

The most common motivation for estimating petrel populations on islands is to inform species status and trend assessments, yet we have shown that the uncertainty around these estimates makes it unlikely that population changes exceeding significant management thresholds could be detected with confidence within typical management timeframes. This suggests that most population estimates cannot reliably inform the management decisions, such as resource allocation and conservation intervention, for which they are intended.

Just over half of the studies we reviewed compared estimates collected at two different times. Of those, most stated that historic estimates were of insufficient quality for a meaningful comparison to be made, with the implicit assertion that the contemporary estimates would now act as a suitable baseline for future comparison. Our results suggest that this implicit assertion may be misplaced. Old sampling designs are often discarded in favour of new ones without adequate thought as to whether the new design is more repeatable and more reliable. The simulation highlighted that after ten years, and under a simplified scenario of no population stochasticity, population declines equivalent to 30% over three generations, the threshold for listing species as Vulnerable on the IUCN Red List, could only be detected in 5% of cases. We stress too that this is merely the detection of a significant difference between two estimates. It does not confirm the magnitude of change that has occurred. In a more realistic scenario of population stochasticity and uncertain variance, even steeper declines would need to occur before a change could be reliably detected. While our simulation was based upon a comparison between estimates at two time points, and it is well known that power could be enhanced by more frequent sampling (Buxton et al., 2016), this is representative of the reality on many islands where petrel surveys may be repeated decades apart using different methodologies. Our findings support the conclusions of focussed single-species or regional studies that current approaches to repeat surveys are likely failing to detect changes and that only very large changes in population size can be detected reliably (Arneill et al., 2019; Hatch, 2003; D. R. Sutherland & Dann, 2012). By simulating population changes equivalent to changes in IUCN Red List status from Least Concern to Vulnerable, Endangered or Critically Endangered it is clear that reported uncertainty in published estimates often precludes detecting changes in conservation status. Criteria-driven assessments e.g. of species status (IUCN, 2012) or of site significance (IUCN, 2016; Ramsar, 2016) have to incorporate uncertainty. While guidelines exist on how this should be done (e.g. IUCN, 2019) they are easier to follow when uncertainty is reported clearly. The level of acceptable uncertainty in population estimates will differ depending on end use, magnitude of acceptable population change and time between surveys. We recommend that prior to implementing new monitoring studies, researchers undertake a power analysis parameterized based on the details of their own system (e.g., species generation length, approximate population size, survey method, magnitude of acceptable decline, timeframe, etc.) to determine levels of uncertainty appropriate to their situation (see Buxton et al., 2016).

### Are population estimates always warranted?

Almost universally the opening sentiments of studies we reviewed highlight the need for “robust”, “accurate”, “precise” population estimates, describing them as “fundamental”, “key”, or “crucial” for conservation, determining trends and informing management (e.g. Arneill et al., 2019; Lavers et al., 2019; Pearson et al., 2013; Sutherland and Dann, 2012; Whitehead et al., 2014). Whether estimates really are warranted depends first on what they are intended for, and second on whether they can achieve the level of certainty required for that stated aim. As is common for biodiversity monitoring in general, estimates were not clearly allied to intended monitoring or management (Possingham et al., 2012). While some studies clearly did inform subsequent management actions either by highlighting unfavourable status and contributing to resulting campaigns (e.g. Cuthbert, 2004) or by providing estimates of potential biological removal to help inform harvest and fisheries management (Defos du Rau et al., 2015; Newman et al., 2009; Rexer-Huber et al., 2017), many did not. Brooke *et al*. (2018) used monitoring data, including population estimates, to derive seabird population trends in response to invasive species eradication. Such syntheses provide an important evidence base for conservation interventions. Similarly, population estimates are useful when they can be linked to clear policy mechanisms such as the Agreement on the Conservation of Albatrosses and Petrels that use population estimates and trends to inform management (Agreement on the Conservation of Albatrosses and Petrels, 2001). However, the collection of estimates because seabirds are ecosystem indicators seems tenuous. Population data for Antarctic Petrels *Fulmaris glaciodes* and Cape Petrels *Daption capensis* can be used in fisheries management by the Convention on the Conservation of Antarctic Marine Living Resources (CCAMLR) Ecosystem Monitoring Program (Agnew, 1997; Descamps et al., 2016), but species captured in our review currently lack equivalent linkages to policy and management.

By clearly articulating their objectives for generating a population estimate, studies can determine the level of acceptable uncertainty (Possingham et al., 2012; Runge et al., 2011). For petrels, survey design will typically be constrained (Arneill et al., 2019), often by resources but also by logistics owing to other variables such as safety requirements or terrain limitations (Oppel et al., 2014). Given these constraints we caution that achievable levels of certainty around population estimates may inevitably be low. It is important that studies do optimise their design given the available resources (both financial and informational), and then carefully and objectively review the potential value of generating a population estimate given the level of uncertainty likely in the result. A significant challenge is that published recommendations are dispersed across a broad literature which can be difficult to access (Poisot et al., 2019). Whereas the emergence of evidence-based conservation as a pillar of conservation science has been supported by projects collating and synthesising evidence to inform management (W. J. Sutherland & Wordley, 2017), no equivalent projects yet collate and synthesise monitoring methods—the basis for the collection of evidence.

Fundamentally, not all reasons for estimating petrel populations demand high levels of precision, but for those that do it is important to objectively assess whether they can be achieved given available resources. If not, resources should either be increased if there is value in doing so, or no attempt should be made to estimate the population.

### Directions for future research

An effective way to build on our *a priori* assessment of uncertainty tolerances would be to conduct a stakeholder-driven value of information analysis to quantify the value of reducing uncertainty in population estimates depending on their intended uses (Canessa et al., 2015; Runge et al., 2011). There is also a need to better understand when and how population estimates can be improved. We found that the rate of publication of petrel population estimates has increased through time. More people are tackling the issue of how to estimate petrel populations, and more research is benefitting from peer review, but it remains a fundamentally challenging system to study. Our meta-analysis suggests uncertainty has actually increased through time despite the emergence of modern technologies and advances in statistical methods (Langrock & Borchers, 2017), although this finding probably reflects poor reporting of early estimates which, if not explicit about uncertainty, were excluded from analysis. While uncertainty may not have been reduced, bias very likely has. Whereas old estimates were more likely to employ biased survey methods like targeted searches, and were less likely to report uncertainty, modern estimates are more likely to be based upon unbiased survey designs, and they are more likely to report uncertainty.

We also found that none of the variables we tested were reliable predictors of the overall level of uncertainty in each study. As a result, there are no easy guidelines around species and geographic traits that can be used at the planning stage to inform decision making around funding and undertaking population estimates, hence the importance of considering whether generating an estimate warrants the effort required.

However, we found that uncertainty was much greater around the estimated number of burrows than it was around burrow occupancy, but more than half the studies did not report uncertainty around their occupancy estimates (Figure 4). This highlights the importance of survey design to minimise variance around burrow density and uncertainty in overall numbers (Appendix S5 in Supporting Information). In terms of occupancy, in some cases, uncertainty may have been estimated and may be factored into the final population estimate, just not reported explicitly in the paper, but in many cases it appears not to have been considered. We recommend sampling occupancy in a number of plots and calculating variance between occupancy rates in all plots so that an overall estimate of the spread of sample means can be generated and confidence intervals reported, or failing that the binomial confidence interval of a proportion is calculated.

We found evidence that mean normalized confidence intervals differed between occupancy methods (Figure S4 and Table S3 in Supporting Information). Other studies allude to the differences in accuracy between different methods of assessing burrow occupancy, but this warrants a quantitative review – something that is only possible if studies report their uncertainty. There is no reason why user-centred design cannot further improve precision around occupancy, ideally eliminating uncertainty through the use of ongoing technological developments.

## Conclusions

Estimates of petrel populations are commonly made to inform trend and threat assessments, but they currently perform rather poorly in this role. Therefore it might be appropriate to focus research on parameters better able to inform management such as productivity and survival (Caravaggi et al., 2019; Dilley et al., 2018) and population trends derived from repeat sampling (Buxton et al., 2016). While population estimates are used in various ways (Table 1) only really when small populations are compared against criteria thresholds for status assessments and when informing 1% thresholds is the actual estimate important. In other cases alternative metrics may be more appropriate. Further methodological improvements could reduce uncertainty, particularly uncertainty around burrow occupancy, but potential gains are finite, and past history suggests that improvement over time is not inevitable or necessarily very rapid. Based on our experience examining historic datasets on multiple species of petrels and undertaking field surveys for contemporary estimates, and now this work, we encourage others to report methods, results and uncertainty clearly and fully to ensure estimates are used appropriately, and to enable further improvements in population estimation. To ensure the best use of conservation funds it is worth considering why a population estimate is being generated, and giving prior consideration to whether the final estimate is likely to achieve a sufficient level of certainty.

## ACKNOWLEDGEMENTS

We would like to thank Simon Wotherspoon, for advice on the analysis, and Holly Jones, the Associate Editor and two anonymous referees for extremely valuable comments and advice for improving the manuscript. JB was supported by a Research Training Program scholarship, an Antarctic Science International Bursary, National Environmental Science Programme Threatened Species Recovery Hub Research Support and a BirdLife Australia Stuart Leslie Bird Research Award. BW was supported by a Postdoctoral Fellowship from the Natural Sciences and Engineering Research Council of Canada and ARC Linkage Project LP150101059 (awarded to RF).

## DATA AVAILABILITY STATEMENT

All data and R code are available from the Australian Antarctic Data Centre: doi:10.4225/15/5282F113C4277.

