## Supporting Information for "Uncertainty in population estimates: a meta-analysis for petrels"

### Appendix S1: Sources of uncertainty in petrel surveys

**Table S1:** Sources of uncertainty in petrel surveys

| Source | Description | Survey adaptation |
| --- | --- | --- |
| Burrow detection | Petrel nests are mostly hidden in underground burrows whose entrances are irregularly distributed, often across challenging terrain and often hidden obscured by dense vegetation (Rayner et al., 2007). Burrow entrances are inactive by day - birds are only active at the colony at night (Warham, 1996). | Counting burrows requires active searching, either of an entire site, or within plots or along transects. Approaches like distance sampling are explicitly designed to account for uncertain detection (Marques et al., 2007). Others may include validation searches to estimate Type II error (e.g. Parker et al., 2015). Aerial surveys using drones can achieve narrow confidence intervals around burrow estimates, but are only effective in sparsely vegetated colonies (Albores-Barajas et al., 2018). |
| Burrow occupancy | Early population estimates typically assumed every burrow entrance represented one breeding attempt, but numerous studies have subsequently highlighted the variability in burrow occupancy and the importance of including an occupancy correction factor in final estimates. Burrows can be long and/or narrow making inspection difficult (Carlile et al., 2019). | Different studies have applied grubbing (feeling with hands or sticks for an occupant; Schulz et al., 2006), recording of signs like feathers, scats, smell etc. (Jahncke and Goya, 1998), playback (Barbraud and Delord, 2006), burrow-scopes (Carlile et al., 2019), and inspection hatches (Cuthbert, 2004) to assess occupancy. Error rates are reported in some cases, and may include ground-truthing – for example playback response rates may be calibrated using a sample of inspection hatches so that the response data can be adjusted (Dilley et al., 2017) – but there has been little comparison of relative error rates achieved by different methods. |
| Measurement error | Any study must balance sampling coverage to build an accurate picture of presence and absence throughout the survey area, sampling intensity to increase confidence around sample means of burrow density and occupancy, and sampling frequency to understand intra- and inter-annual variation in occupancy. | In practice this balance is often dictated by costs and logistics (Arneill et al., 2019). Sampling of multiple sites across the density gradient allows for calculation of confidence intervals around the sample mean, but owing to high variability in burrow density, and zero-inflated data, uncertainty can remain high until the sample size becomes very large (Sileshi et al., 2009). |
| Temporal variation | Repeat sampling is important for understanding temporal variation but is not always possible if the study site can only be accessed infrequently. | Emerging approaches such as camera traps to monitor burrow occupancy and breeding status of burrows may allow measurement of temporal variation (Bird et al., submitted). |

### Appendix S2: species included in the WoS search

We searched the Web of Science bibliographic index on 20 January 2020 using the search terms "burrowing seabird" OR "burrow-nesting seabird" OR "burrow-nesting petrel" OR "burrowing petrel" OR “scientific name” OR “common name” (taxonomy followed HBW & BirdLife International, 2018) for all species in the families *Procellariidae*, *Hydrobatidae* and *Oceanitidae*, AND “abundance” OR “population” in the title, abstract or keywords.

**Table S2:** Of the 124 petrel species searches were restricted to 110 burrow/crevice/cavity nesting species

| Scientific name | Common name | Scientific name | Common name |
|---|---|---|---|
| Oceanites oceanicus | Wilson's Storm-petrel | Pterodroma hasitata | Black-capped Petrel |
| Oceanites gracilis | White-vented Storm-petrel | Pterodroma caribbaea | Jamaican Petrel |
| Oceanites pincoyae | Pincoya Storm-petrel | Pterodroma feae | Cape Verde Petrel |
| Garrodia nereis | Grey-backed Storm-petrel | Pterodroma deserta | Desertas Petrel |
| Pelagodroma marina | White-faced Storm-petrel | Pterodroma madeira | Zino's Petrel |
| Fregetta grallaria | White-bellied Storm-petrel | Pterodroma magentae | Magenta Petrel |
| Fregetta tropica | Black-bellied Storm-petrel | Pterodroma incerta | Atlantic Petrel |
| Fregetta maoriana | New Zealand Storm-petrel | Pterodroma lessonii | White-headed Petrel |
| Nesofregetta fuliginosa | Polynesian Storm-petrel | Pterodroma macroptera | Great-winged Petrel |
| Hydrobates pelagicus | European Storm-petrel | Pterodroma gouldi | Grey-faced Petrel |
| Hydrobates jabejabe | Cape Verde Storm-petrel | Procellaria cinerea | Grey Petrel |
| Hydrobates castro | Band-rumped Storm-petrel | Procellaria aequinoctialis | White-chinned Petrel |
| Hydrobates monteiroi | Monteiro's Storm-petrel | Procellaria conspicillata | Spectacled Petrel |
| Hydrobates matsudairae | Matsudaira's Storm-petrel | Procellaria westlandica | Westland Petrel |
| Hydrobates melania | Black Storm-petrel | Procellaria parkinsoni | Black Petrel |
| Hydrobates homochroa | Ashy Storm-petrel | Ardenna pacifica | Wedge-tailed Shearwater |
| Hydrobates microsoma | Least Storm-petrel | Ardenna bulleri | Buller's Shearwater |
| Hydrobates tethys | Wedge-rumped Storm-petrel | Ardenna tenuirostris | Short-tailed Shearwater |
| Hydrobates socorroensis | Townsend's Storm-petrel | Ardenna grisea | Sooty Shearwater |
| Hydrobates cheimomnestes | Ainley's Storm-petrel | Ardenna gravis | Great Shearwater |
| Hydrobates leucorhous | Leach's Storm-petrel | Ardenna carneipes | Flesh-footed Shearwater |
| Hydrobates monorhis | Swinhoe's Storm-petrel | Ardenna creatopus | Pink-footed Shearwater |
| Hydrobates macrodactylus | Guadalupe Storm-petrel | Calonectris leucomelas | Streaked Shearwater |
| Hydrobates tristrami | Tristram's Storm-petrel | Calonectris diomedea | Scopoli's Shearwater |
| Hydrobates markhami | Markham's Storm-petrel | Calonectris borealis | Cory's Shearwater |
| Hydrobates furcatus | Fork-tailed Storm-petrel | Calonectris edwardsii | Cape Verde Shearwater |
| Hydrobates hornbyi | Ringed Storm-petrel | Puffinus subalaris | Galapagos Shearwater |
| Pagodroma nivea | Snow Petrel | Puffinus gavia | Fluttering Shearwater |
| Halobaena caerulea | Blue Petrel | Puffinus huttoni | Hutton's Shearwater |
| Pachyptila vittata | Broad-billed Prion | Puffinus opisthomelas | Black-vented Shearwater |
| Pachyptila salvini | Salvin's Prion | Puffinus bryani | Bryan's Shearwater |
| Pachyptila macgillivrayi | MacGillivray's Prion | Puffinus myrtae | Rapa Shearwater |
| Pachyptila desolata | Antarctic Prion | Puffinus newelli | Newell's Shearwater |

| Scientific name | Common name | Scientific name | Common name |
|---|---|---|---|
| Pachyptila belcheri | Slender-billed Prion | Puffinus auricularis | Townsend's Shearwater |
| Pachyptila turtur | Fairy Prion | Puffinus bailloni | Tropical Shearwater |
| Pachyptila crassirostris | Fulmar Prion | Puffinus persicus | Persian Shearwater |
| Aphrodroma brevirostris | Kerguelen Petrel | Puffinus bannermani | Bannerman's Shearwater |
| Pterodroma rupinarum | Large St Helena Petrel | Puffinus puffinus | Manx Shearwater |
| Pterodroma leucoptera | White-winged Petrel | Puffinus yelkouan | Yelkouan Shearwater |
| Pterodroma brevipes | Collared Petrel | Puffinus mauretanicus | Balearic Shearwater |
| Pterodroma defilippiana | Masatierra Petrel | Puffinus elegans | Subantarctic Shearwater |
| Pterodroma longirostris | Stejneger's Petrel | Puffinus assimilis | Little Shearwater |
| Pterodroma cookii | Cook's Petrel | Puffinus lherminieri | Audubon's Shearwater |
| Pterodroma pycrofti | Pycroft's Petrel | Puffinus heinrothi | Heinroth's Shearwater |
| Pterodroma hypoleuca | Bonin Petrel | Pseudobulweria macgillivrayi | Fiji Petrel |
| Pterodroma nigripennis | Black-winged Petrel | Pseudobulweria aterrima | Mascarene Petrel |
| Pterodroma axillaris | Chatham Petrel | Pseudobulweria becki | Beck's Petrel |
| Pterodroma baraui | Barau's Petrel | Pseudobulweria rostrata | Tahiti Petrel |
| Pterodroma inexpectata | Mottled Petrel | Bulweria bulwerii | Bulwer's Petrel |
| Pterodroma sandwichensis | Hawaiian Petrel | Bulweria fallax | Jouanin's Petrel |
| Pterodroma phaeopygia | Galapagos Petrel | Bulweria bifax | Small St Helena Petrel |
| Pterodroma cervicalis | White-necked Petrel | Pelecanoides garnotii | Peruvian Diving-petrel |
| Pterodroma externa | Juan Fernandez Petrel | Pelecanoides magellani | Magellanic Diving-petrel |
| Pterodroma mollis | Soft-plumaged Petrel | Pelecanoides georgicus | South Georgia Diving-petrel |
| Pterodroma cahow | Bermuda Petrel | Pelecanoides urinatrix | Common Diving-petrel |

### Appendix S3: Simulated power analysis

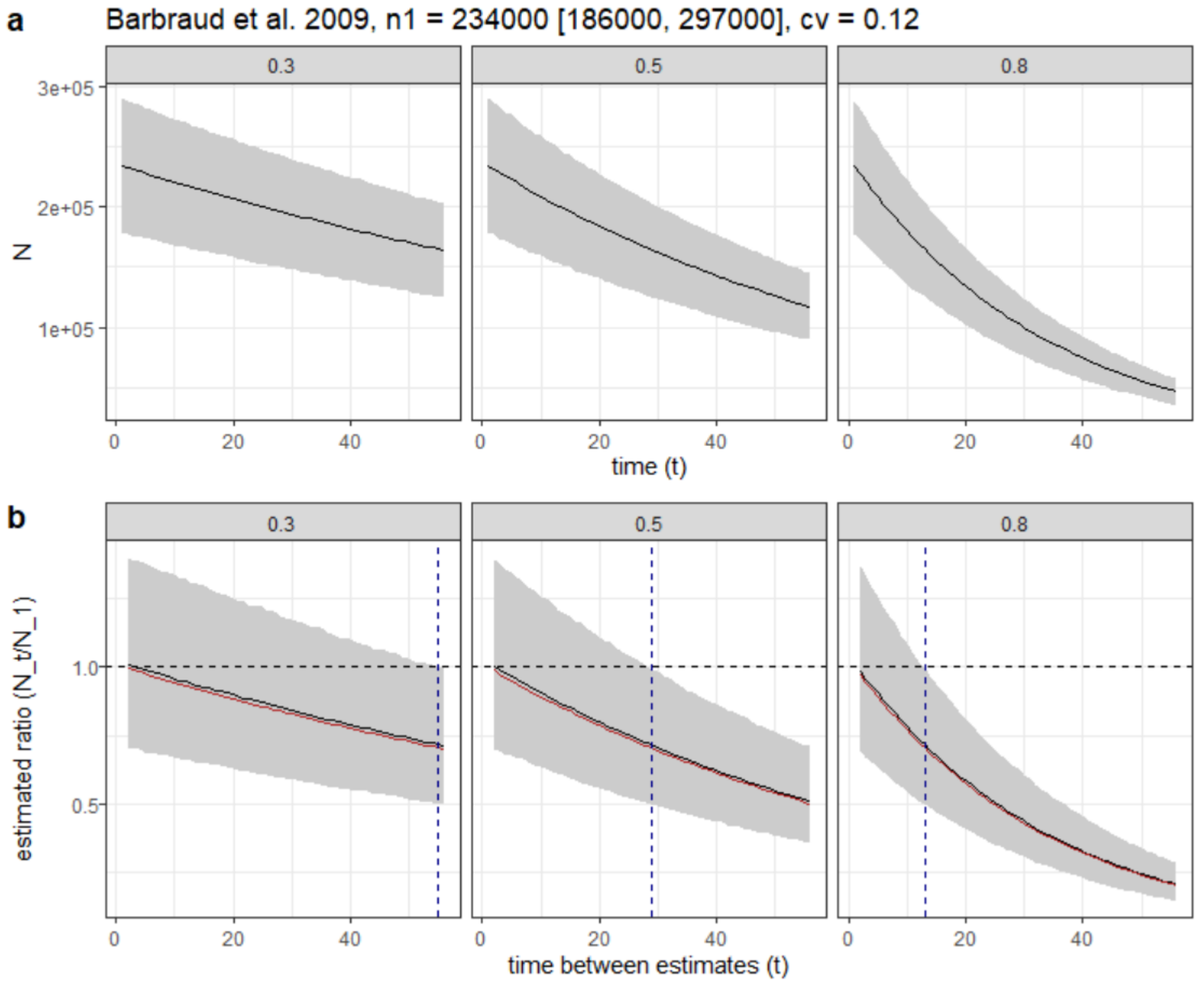


**Figure S1: (a)** Simulated population declines of 30%, 50% and 80% over three generations; (b) Proportion of Monte Carlo simulations where differences in mean estimates indicated a decline, N_t/N_1<1 (solid line). **(b)** Mean estimated rate of decline (N_t/N_1) with 95% CI over different sampling intervals. The horizontal black dashed line represents no change (N_t/N_1=1), and the vertical dashed line shows the first time-step at which the upper 95% CI excludes 1.

#### Appendix S4: Publishing trends and correlates of uncertainty

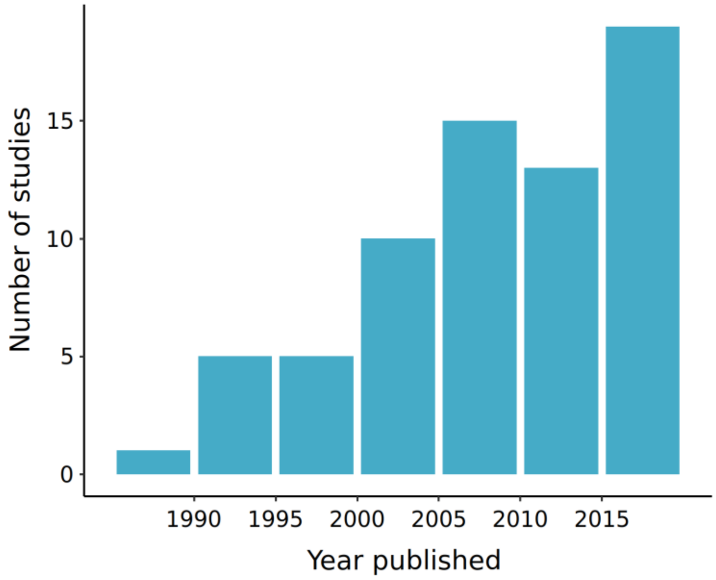


**Figure S2:** Number of studies publishing petrel population estimates through time

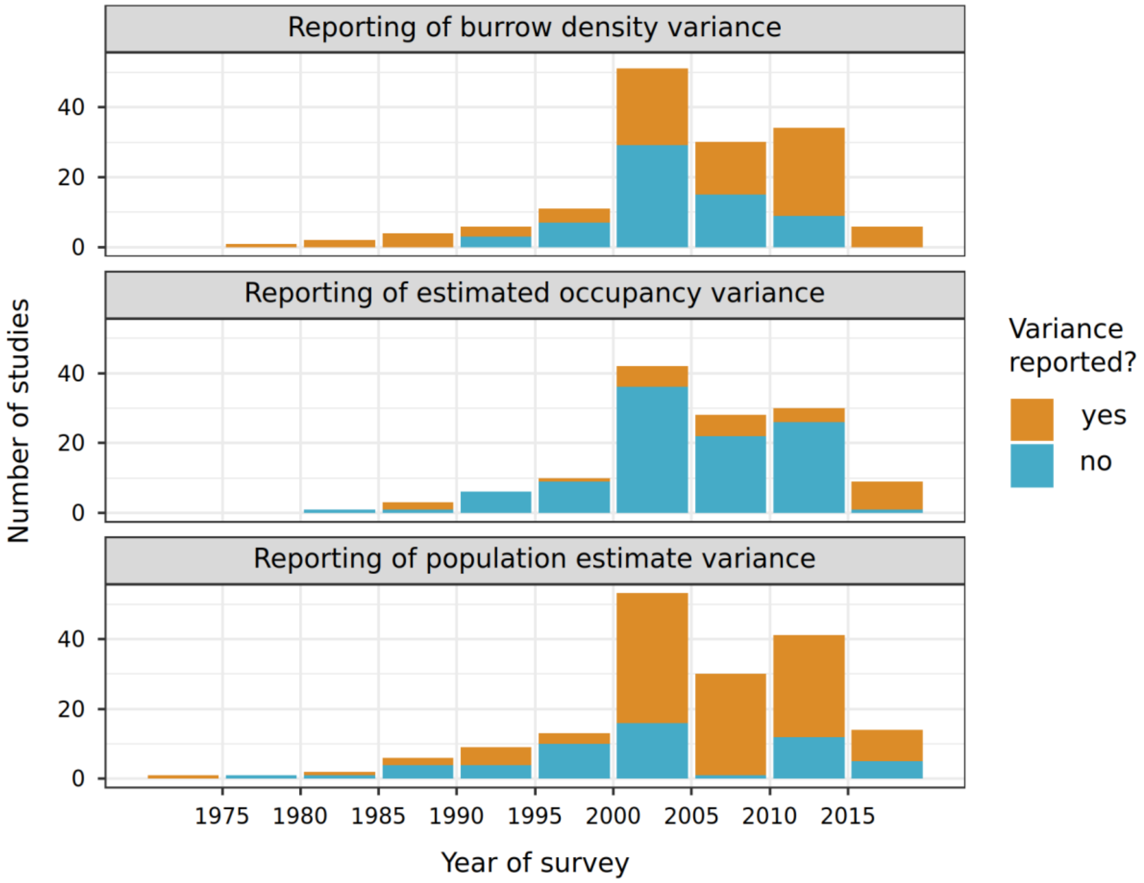


**Figure S3:** Number of studies that reported variance with published estimates of burrow density, occupancy, and population size over time.

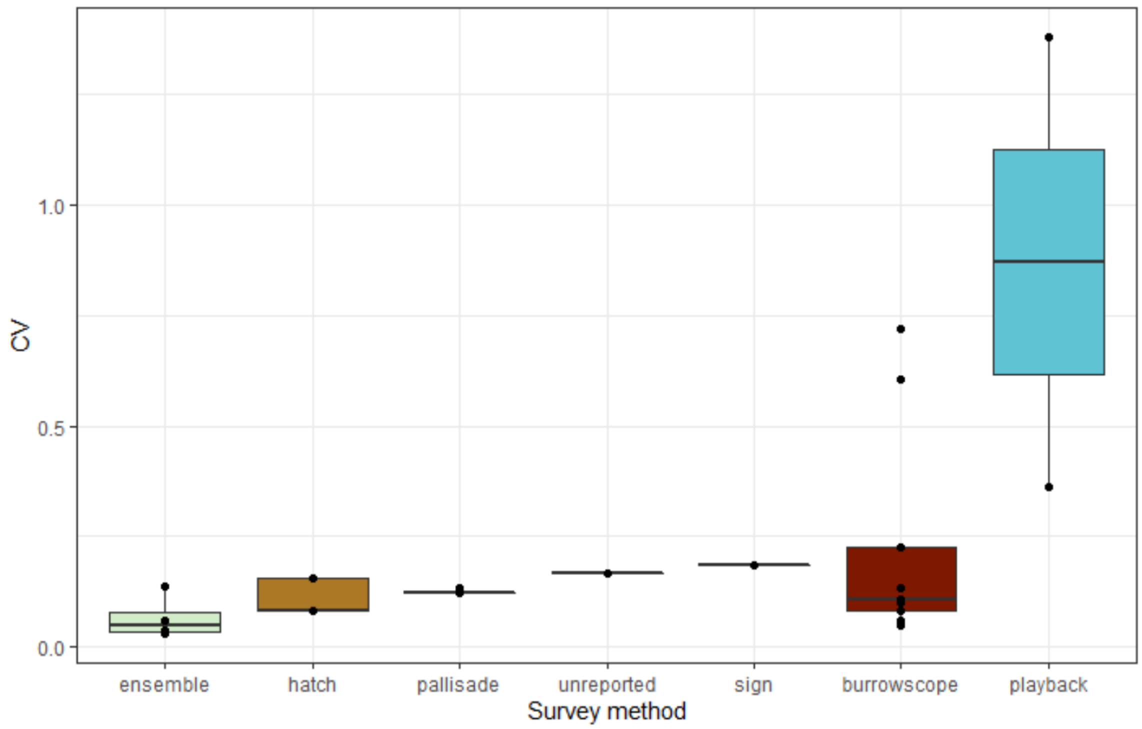


**Figure S4:** Uncertainty in estimates of burrow occupancy based upon different survey methods

**Table S3:** Analysis of Variance and Tukey's multiple comparisons of means suggest uncertainty in burrow occupancy when using playback is significantly higher than when using hatches, or an ensemble method combining multiple approaches.

| ANOVA | | | | | |
|---|---|---|---|---|---|
| | Df | Sum Sq | Mean Sq | F value | P value |
| | 5, 20 | 9.2143 | 1.84285 | 3.6039 | 0.01739 |

Tukey multiple comparisons of means:

Fit: aov(formula = log(occ_cv) ~ 1 + occ_method, data = .)

| | diff | lwr | upr | P adj |
|---|---|---|---|---|
| hatch-burrowscope | -0.34182 | -1.59553 | 0.911898 | 0.952414 |
| pallisade-burrowscope | -0.16088 | -1.4146 | 1.092832 | 0.998406 |
| ensemble-burrowscope | -0.9844 | -2.3351 | 0.36631 | 0.243213 |
| playback-burrowscope | 1.571121 | -0.186 | 3.328238 | 0.096716 |
| sign-burrowscope | 0.229083 | -2.14021 | 2.59838 | 0.999594 |
| pallisade-hatch | 0.180934 | -1.24064 | 1.602512 | 0.998467 |
| ensemble-hatch | -0.64258 | -2.15039 | 0.865231 | 0.760382 |
| playback-hatch | 1.912937 | 0.032366 | 3.793508 | 0.044744 |
| sign-hatch | 0.570899 | -1.89135 | 3.033145 | 0.975924 |
| ensemble-pallisade | -0.82351 | -2.33133 | 0.684297 | 0.537117 |
| playback-pallisade | 1.732003 | -0.14857 | 3.612574 | 0.082157 |
| sign-pallisade | 0.389965 | -2.07228 | 2.852211 | 0.995688 |
| playback-ensemble | 2.555518 | 0.608941 | 4.502094 | 0.005994 |
| sign-ensemble | 1.21348 | -1.29954 | 3.726499 | 0.657503 |
| sign-playback | -1.34204 | -4.09491 | 1.410837 | 0.648766 |

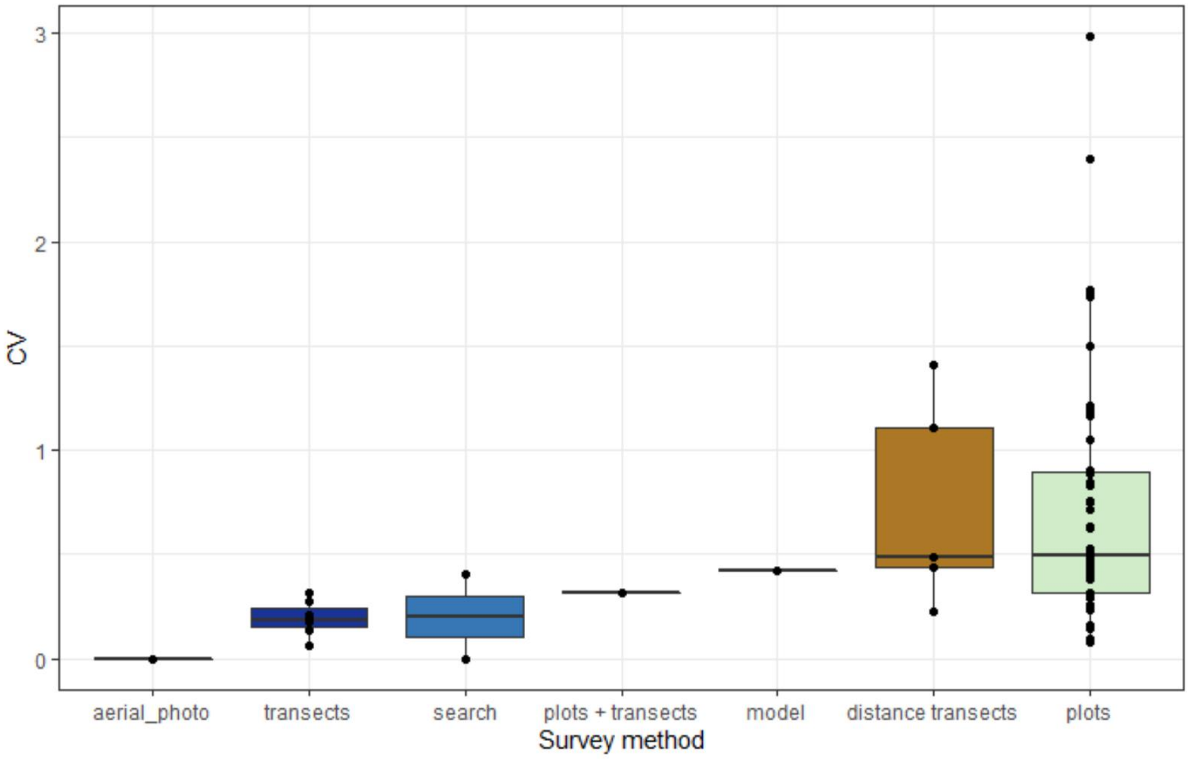


**Figure S5:** Uncertainty in estimates of burrow numbers based upon different survey methods

**Table S3:** Analysis of Variance and Tukey's multiple comparisons of means suggest uncertainty in burrow numbers when using transects is significantly lower than when using plots, but we found no difference between either method and distance transects.

| ANOVA | | | | | |
|---|---|---|---|---|---|
| | Df | Sum Sq | Mean Sq | F value | P value |
| | 5, 56 | 8.465 | 1.693 | 2.7882 | 0.02561 |
| Tukey multiple comparisons of means: | | | | | |
| Fit: aov(formula = log(brw_cv) ~ 1 + brw_method, data = .) | | | | | |
| | diff | lwr | upr | P adj | |
| model-distance transects | -0.3497 | -2.2734 | 1.574006 | 0.99441 | |
| plots-distance transects | -0.08463 | -1.16734 | 0.998074 | 0.999906 | |
| plots + transects-distance transects | -0.63818 | -3.1569 | 1.880533 | 0.974876 | |
| search-distance transects | -0.3922 | -2.91092 | 2.126513 | 0.997303 | |
| transects-distance transects | -1.23049 | -2.5768 | 0.115823 | 0.091922 | |
| plots-model | 0.265067 | -1.39573 | 1.92586 | 0.996968 | |
| plots + transects-model | -0.28849 | -3.1045 | 2.527524 | 0.999643 | |
| search-model | -0.04251 | -2.85852 | 2.773504 | 1 | |
| transects-model | -0.88079 | -2.7243 | 0.962722 | 0.721019 | |
| plots + transects-plots | -0.55355 | -2.87768 | 1.770567 | 0.980858 | |
| search-plots | -0.30757 | -2.6317 | 2.016547 | 0.998762 | |
| transects-plots | -1.14586 | -2.07868 | -0.21304 | 0.007834 | |
| search-plots + transects | 0.24598 | -3.00567 | 3.497631 | 0.99992 | |
| transects-plots + transects | -0.5923 | -3.05032 | 1.865714 | 0.979846 | |
| transects-search | -0.83828 | -3.2963 | 1.619734 | 0.91393 | |

**Table S4:** Top 10 multi-variate models of CV related to explanatory variables. The best model is an intercept only model suggesting no explanatory variables explain uncertainty in population estimates

| (Intercept) | method | rl_status | year_est | zbody_mass | zisland_area | zpop_est | df | logLik | AICc | delta | weight |
|---|---|---|---|---|---|---|---|---|---|---|---|
| 0.165101 | NA | NA | NA | NA | NA | NA | 3 | 57.74026 | -109.231 | 0 | 0.956245 |
| 0.165663 | NA | NA | NA | -0.00807 | NA | NA | 4 | 54.6048 | -100.789 | 8.44196 | 0.014042 |
| 0.164755 | NA | NA | NA | NA | 0.004319 | NA | 4 | 54.47825 | -100.535 | 8.695059 | 0.012373 |
| 0.165349 | NA | NA | NA | NA | NA | -0.00388 | 4 | 54.41792 | -100.415 | 8.815727 | 0.011648 |
| -5.4141 | NA | NA | 0.002782 | NA | NA | NA | 4 | 53.49537 | -98.5697 | 10.66083 | 0.00463 |
| 0.118972 | NA | + | NA | NA | NA | NA | 6 | 52.98607 | -93.0689 | 16.1616 | 0.000296 |
| 0.165308 | NA | NA | NA | -0.01143 | 0.00859 | NA | 5 | 51.52757 | -92.4168 | 16.81366 | 0.000214 |
| 0.16584 | NA | NA | NA | -0.00775 | NA | -0.00324 | 5 | 51.27786 | -91.9174 | 17.3131 | 0.000166 |
| 0.164999 | NA | NA | NA | NA | 0.004383 | -0.00389 | 5 | 51.16159 | -91.6849 | 17.54564 | 0.000148 |
| -6.05961 | NA | NA | 0.003104 | -0.0133 | NA | NA | 5 | 50.5893 | -90.5403 | 18.69022 | 8.36E-05 |

**Table S5:** Tukey HSD multiple comparisons of mean CV of population estimates based upon different survey methods. Survey methods are derived from Rayner et al. (2007): mark recapture (MR), counts of known colonies (C[KL]), simple extrapolation (S[S]), simple habitat area models (S[HA]), advanced habitat area models (A[HA]), informed estimates (IE).

| comparison | diff | lwr | upr | p adj |
|---|---|---|---|---|
| C(KL)-A(HA) | -0.80573 | -1.80996 | 0.198507 | 0.177423 |
| MR-A(HA) | -0.33375 | -1.70229 | 1.034787 | 0.960763 |
| S(HA)-A(HA) | -0.21456 | -1.06571 | 0.636591 | 0.955859 |
| S(S)-A(HA) | -0.276 | -1.11251 | 0.560522 | 0.889664 |
| MR-C(KL) | 0.471974 | -0.84287 | 1.786823 | 0.855744 |
| S(HA)-C(KL) | 0.591164 | -0.17067 | 1.352998 | 0.205028 |
| S(S)-C(KL) | 0.529729 | -0.21572 | 1.275178 | 0.285804 |
| S(HA)-MR | 0.119189 | -1.08281 | 1.32119 | 0.998713 |
| S(S)-MR | 0.057755 | -1.13393 | 1.249438 | 0.999925 |
| S(S)-S(HA) | -0.06143 | -0.58266 | 0.459788 | 0.99747 |

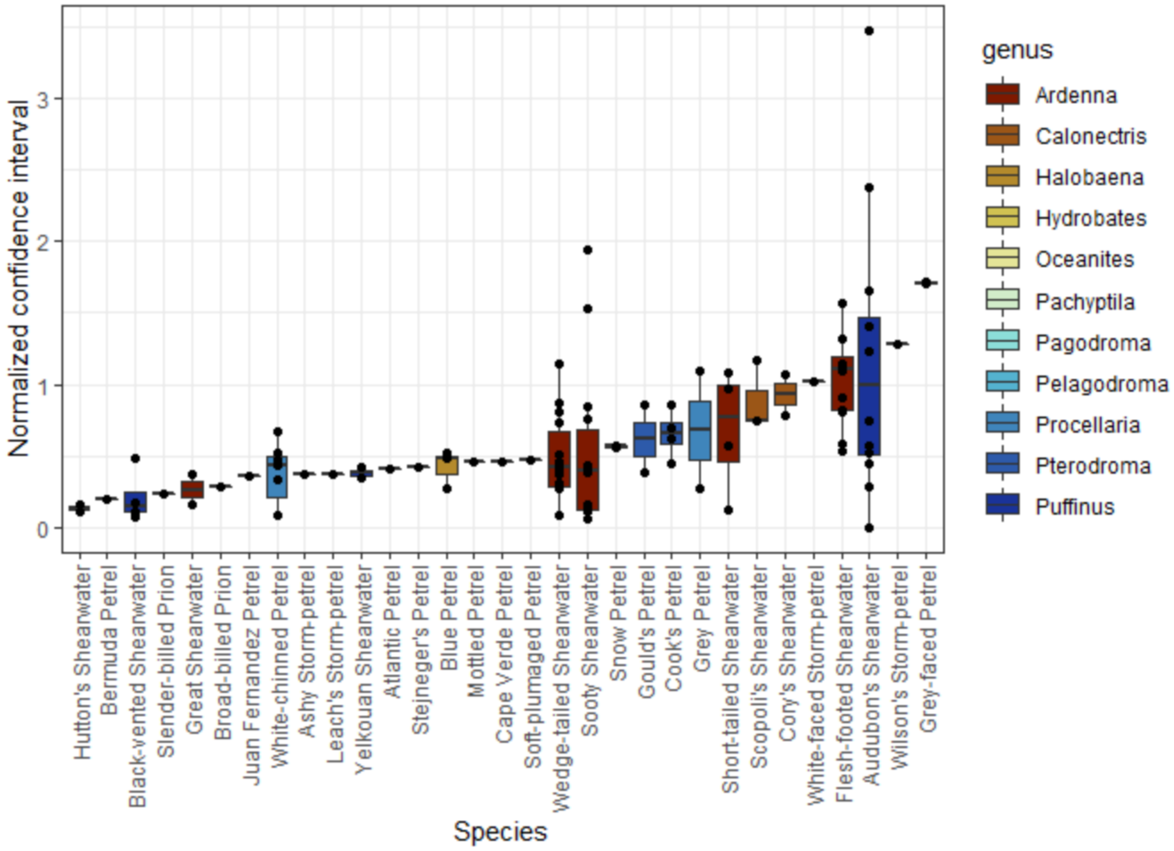


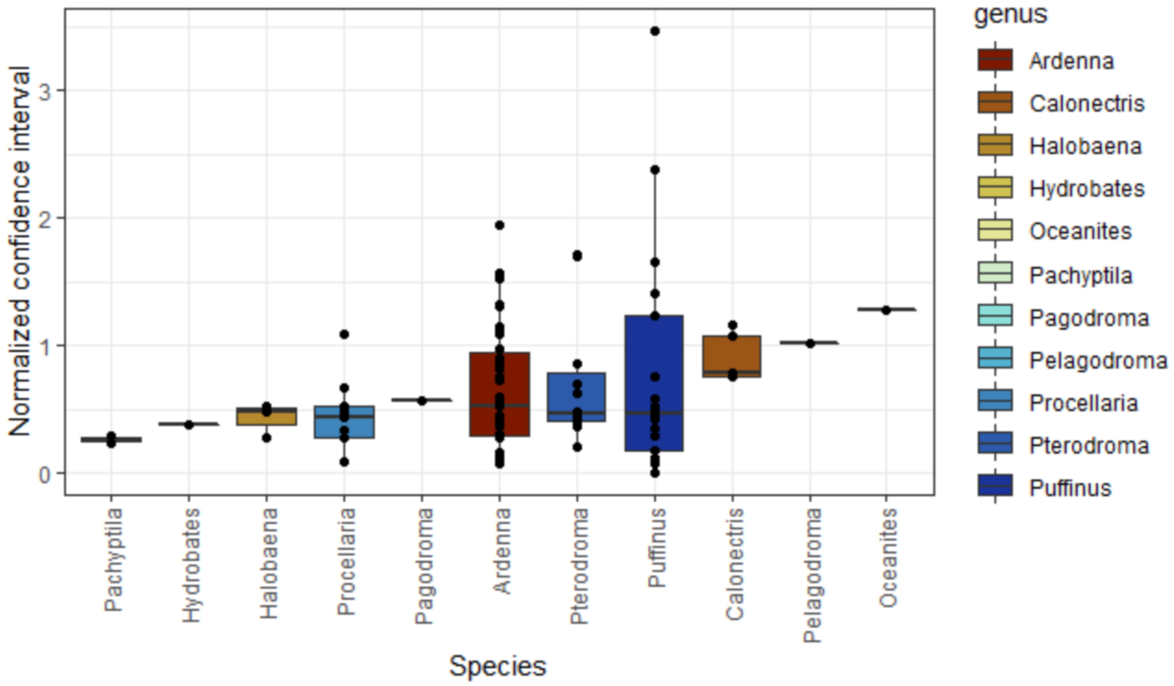


**Figure S6:** Uncertainty in population estimates by species and genus

**Table S2:** Analysis of Variance and Tukey's multiple comparisons of means provided no evidence of a difference in uncertainty in population estimates between genera.

| ANOVA | | | | | |
|---|---|---|---|---|---|
| Df | | Sum Sq | Mean Sq | F value | P value |
| genus | 10 | 2.2779 | 0.22779 | 0.7591 | 0.6673 |

Tukey multiple comparisons of means:

Fit: aov(formula = ci_norm ~ 1 + genus, data = .)

| | diff | lwr | upr | P adj |
|---|---|---|---|---|
| Calonectris-Ardenna | 0.256334 | -0.59451 | 1.107178 | 0.995533 |
| Halobaena-Ardenna | -0.2241 | -1.30078 | 0.85258 | 0.999813 |
| Hydrobates-Ardenna | -0.27475 | -1.57988 | 1.030388 | 0.999793 |
| Oceanites-Ardenna | 0.630372 | -1.19604 | 2.456785 | 0.987146 |
| Pachyptila-Ardenna | -0.39121 | -1.69634 | 0.913927 | 0.995711 |
| Pagodroma-Ardenna | -0.08307 | -1.3882 | 1.22207 | 1 |
| Pelagodroma-Ardenna | 0.367517 | -1.4589 | 2.19393 | 0.999862 |
| Procellaria-Ardenna | -0.20984 | -0.86842 | 0.448745 | 0.993034 |
| Pterodroma-Ardenna | 0.021827 | -0.51541 | 0.559068 | 1 |
| Puffinus-Ardenna | 0.143322 | -0.34064 | 0.627279 | 0.99611 |
| Halobaena-Calonectris | -0.48044 | -1.79999 | 0.839121 | 0.980941 |
| Hydrobates-Calonectris | -0.53108 | -2.04282 | 0.980662 | 0.985334 |
| Oceanites-Calonectris | 0.374038 | -1.6053 | 2.353374 | 0.999922 |
| Pachyptila-Calonectris | -0.64754 | -2.15929 | 0.8642 | 0.942223 |
| Pagodroma-Calonectris | -0.3394 | -1.85114 | 1.172344 | 0.999632 |
| Pelagodroma-Calonectris | 0.111183 | -1.86815 | 2.090519 | 1 |
| Procellaria-Calonectris | -0.46617 | -1.474 | 0.541657 | 0.907447 |
| Pterodroma-Calonectris | -0.23451 | -1.16758 | 0.698561 | 0.999013 |
| Puffinus-Calonectris | -0.11301 | -1.01645 | 0.790427 | 0.999998 |
| Hydrobates-Halobaena | -0.05064 | -1.70009 | 1.598802 | 1 |
| Oceanites-Halobaena | 0.854475 | -1.23193 | 2.940878 | 0.956844 |
| Pachyptila-Halobaena | -0.16711 | -1.81655 | 1.48234 | 1 |
| Pagodroma-Halobaena | 0.141037 | -1.50841 | 1.790484 | 1 |
| Pelagodroma-Halobaena | 0.591619 | -1.49478 | 2.678023 | 0.997272 |
| Procellaria-Halobaena | 0.014265 | -1.19032 | 1.21885 | 1 |
| Pterodroma-Halobaena | 0.245929 | -0.89684 | 1.388699 | 0.999747 |
| Puffinus-Halobaena | 0.367424 | -0.75129 | 1.486134 | 0.991184 |
| Oceanites-Hydrobates | 0.905119 | -1.30785 | 3.118085 | 0.957218 |
| Pachyptila-Hydrobates | -0.11646 | -1.92334 | 1.690417 | 1 |
| Pagodroma-Hydrobates | 0.191682 | -1.6152 | 1.99856 | 1 |
| Pelagodroma-Hydrobates | 0.642264 | -1.5707 | 2.855229 | 0.996699 |
| Procellaria-Hydrobates | 0.06491 | -1.34759 | 1.477412 | 1 |
| Pterodroma-Hydrobates | 0.296574 | -1.06359 | 1.656743 | 0.999716 |
| Puffinus-Hydrobates | 0.418069 | -0.92195 | 1.758086 | 0.994097 |
| Pachyptila-Oceanites | -1.02158 | -3.23455 | 1.191384 | 0.908522 |
| Pagodroma-Oceanites | -0.71344 | -2.9264 | 1.499528 | 0.992365 |
| Pelagodroma-Oceanites | -0.26286 | -2.81817 | 2.292457 | 1 |
| Procellaria-Oceanites | -0.84021 | -2.74483 | 1.064407 | 0.930485 |
| Pterodroma-Oceanites | -0.60855 | -2.47468 | 1.257591 | 0.991656 |
| Puffinus-Oceanites | -0.48705 | -2.33855 | 1.364449 | 0.998543 |

| | | | | |
|---|---|---|---|---|
| Pagodroma-Pachyptila | 0.308144 | -1.49873 | 2.115022 | 0.99997 |
| Pelagodroma-Pachyptila | 0.758726 | -1.45424 | 2.971691 | 0.987772 |
| Procellaria-Pachyptila | 0.181372 | -1.23113 | 1.593873 | 0.999998 |
| Pterodroma-Pachyptila | 0.413036 | -0.94713 | 1.773204 | 0.99524 |
| Puffinus-Pachyptila | 0.534531 | -0.80549 | 1.874548 | 0.96383 |
| Pelagodroma-Pagodroma | 0.450582 | -1.76238 | 2.663547 | 0.999847 |
| Procellaria-Pagodroma | -0.12677 | -1.53927 | 1.28573 | 1 |
| Pterodroma-Pagodroma | 0.104892 | -1.25528 | 1.465061 | 1 |
| Puffinus-Pagodroma | 0.226387 | -1.11363 | 1.566404 | 0.999973 |
| Procellaria-Pelagodroma | -0.57735 | -2.48197 | 1.327263 | 0.995306 |
| Pterodroma-Pelagodroma | -0.34569 | -2.21183 | 1.520446 | 0.999935 |
| Puffinus-Pelagodroma | -0.22419 | -2.07569 | 1.627305 | 0.999999 |
| Pterodroma-Procellaria | 0.231664 | -0.53018 | 0.993511 | 0.995187 |
| Puffinus-Procellaria | 0.353159 | -0.3721 | 1.078416 | 0.875838 |
| Puffinus-Pterodroma | 0.121495 | -0.49567 | 0.738662 | 0.999887 |

### Appendix S5: Discussion of considerations for selecting survey methods

There are now a number of studies that offer valuable advice on how to design surveys to minimise uncertainty around population estimates and to increase power to detect trends (Arneill et al., 2019; Buxton et al., 2016; Hatch, 2003; McKechnie et al., 2009; Schumann et al., 2013; Sutherland and Dann, 2012). Most studies arrive at petrel population estimates by combining measures of burrow density, burrow occupancy, and the area occupied by burrows. Therefore, to detect overall population changes a method must estimate each of these variables. Arneill et al. (2019) show that the most efficient way to do this is to use a multi-staged stratified sampling approach and to subjectively distribute monitoring plots in high-density areas. This approach can fail to detect population expansion because monitoring plots may be at or close to carrying capacity (Arneill et al., 2019; Jackson et al., 2008). However, it is rare for management criteria to be triggered by threshold increases in seabird populations, so the practical implications of this limitation are low. Population declines will either be driven by i) factors independent of colony density such as bycatch, in which case this sampling approach will still be effective; ii) factors exacerbated by high density such as communicable diseases or invasive species in which case declines will also still be captured by the sampling approach; or by iii) factors exacerbated by low density in which case declines may be undetected or underestimated but the magnitude of this error is minimised by the sampling approach because high density plots support a high proportion of the overall population.

Natural inter-annual variability in burrow occupancy in colonies can be high (Hatch, 2003). Understanding this variability via repeat sampling across years is essential if surveys are to detect trends outside the range of natural fluctuations. Although repeat surveying may be beyond the resources of many projects currently (Arneill et al., 2019) this requires that projects be better resourced, rather than simply going ahead when destined to fail. The advantages of this are two-fold. As well as providing the statistical rigour to ensure population estimates are fit-for-purpose, it also has benefits for the conservation workforce. Job security is particularly low for ecology graduates (Hance, 2017). Many workers are employed on seasonal contracts or simply volunteer to participate in surveys. By planning repeat annual surveys in multi-year pulses it may be possible to offer a greater degree of security to workers than is provided by a one-off survey.

It is common to use prior knowledge to help identify colonies and estimate abundance within them. There are advantages to this targeted approach in terms of survey efficiency and minimising uncertainty (Arneill et al., 2019; Dilley et al., 2019). A drawback is that uncertainty about the proportion of the population missed in a survey because it lies outside the identified focal area is unknown, unless a targeted survey is linked to a randomised survey (Dilley et al., 2019).

Other sources of error relate to temporal variation in burrow occupancy, errors in estimates of occupied area when extrapolating from sample data, and error in identifying burrow occupants to species level. Of the studies we reviewed, six estimated target populations over multiple years and incorporated inter-annual variability in their final population estimate. Not accounting for inter-annual variation of this nature can mask population trends (Hatch, 2003).

Reporting of variance is inconsistent (e.g. McDonald, 2014). Several of the studies we reviewed failed to clarify whether reported variances were standard errors or confidence intervals. When variance in burrow density and burrow occupancy was measured it was unclear how the variance of their products was calculated in the final population estimate. For example, whether burrow density and occupancy estimates were from independent samples or if they were measured in the same sample with variance in apparently occupied burrows within each sampling unit extrapolated for the final population estimate (as in: Arneill et al., 2019; Pearson et al., 2013).

Finally, if setting out to assess trends, it is important to consider uncertainty of historic estimates and to determine the power that can be achieved given available resources (Buxton et al., 2016; Hatch, 2003).
